# MetagenomicKG: a knowledge graph for metagenomic applications

**DOI:** 10.1101/2024.03.14.585056

**Authors:** Chunyu Ma, Shaopeng Liu, David Koslicki

## Abstract

**Motivation:** The sheer volume and variety of genomic content within microbial communities makes metagenomics a field rich in biomedical knowledge. To traverse these complex communities and their vast unknowns, metagenomic studies often depend on distinct reference databases, such as the Genome Taxonomy Database (GTDB), the Kyoto Encyclopedia of Genes and Genomes (KEGG), and the Bacterial and Viral Bioinformatics Resource Center (BV-BRC), for various analytical purposes. These databases are crucial for genetic and functional annotation of microbial communities. Nevertheless, the inconsistent nomenclature or identifiers of these databases present challenges for effective integration, representation, and utilization. Knowledge graphs (KGs) offer an appropriate solution by organizing biological entities and their interrelations into a cohesive network. The graph structure not only facilitates the unveiling of hidden patterns but also enriches our biological understanding with deeper insights. Despite KGs having shown potential in various biomedical fields, their application in metagenomics remains underexplored.

**Results:** We present `MetagenomicKG`, a novel knowledge graph specifically tailored for metagenomic analysis. `MetagenomicKG` integrates taxonomic, functional, and pathogenesis-related information from widely used databases, and further links these with established biomedical knowledge graphs to expand biological connections. Through several use cases, we demonstrate its utility in enabling hypothesis generation regarding the relationships between microbes and diseases, generating sample-specific graph embeddings, and providing robust pathogen prediction.

**Availability and Implementation:** The source code and technical details for constructing the `MetagenomicKG` and reproducing all analyses are available at Github: https://github.com/KoslickiLab/MetagenomicKG. We also host a Neo4j instance: http://mkg.cse.psu.edu:7474 for accessing and querying this graph.

**Supplementary information:** available at *Bioinformatics* online.

## 1 Introduction

Metagenomics, the study of microbial genetic material recovered directly from environmental samples, has unveiled the vast diversity of microbial communities and their roles (Ko *et al*., 2022; Shu and Huang, 2022; Mendes *et al*., 2017). This field offers profound insights into the composition and function of microbial communities and aids the discovery of novel genes, enzymes, and metabolic pathways applicable in medicine, biotechnology, and environmental science. The broad spectrum of microbial life also suggests a significant volume of unknowns, commonly referred to as “microbial dark matter” (Jiao *et al*., 2021; Bodor *et al*., 2020; Pedrós-Alió and Manrubia, 2016). These unknowns include genetic material from unknown and uncultured microbes, along with unknown genetic functions and interactions within microbial communities and their environments. It is thus important for researchers to distinguish between what is genuinely novel and what has been previously observed or analyzed. Therefore, the metagenomic field inherently relies on extensive and comprehensive reference databases to decipher microbial communities and to push the boundaries of knowledge. Numerous well-established reference databases are pivotal in metagenomic studies. The NCBI GenBank (Benson *et al*., 2012) and the Genome Taxonomy Database (GTDB) (Parks *et al*., 2022) databases offer a taxonomic framework for bacterial, archaeal and viral genomes, facilitating a better understanding of microbial phylogeny and taxonomy. The Kyoto Encyclopedia of Genes and Genomes (KEGG) (Kanehisa *et al*., 2017) is a comprehensive source of information on genomes, biological pathways, diseases, and pharmaceuticals, which are crucial for functional annotation. The Bacterial and Viral Bioinformatics Resource Center (BV-BRC) (Olson *et al*., 2023) specializes in bacterial pathogens, offering an array of data and tools for comparative genomic analysis to detect and characterize pathogens. Collectively, these databases and others form the backbone of metagenomic studies, enabling researchers to annotate genetic sequences, decipher microbial functions, and delve into the genetic basis of microbial communities. Furthermore, extensive knowledge about microbes and non-microbes has been established in public databases (Liu *et al*., 2022; Lakin et al., 2017; Consortium, 2010). Nevertheless, although these databases are inherently linked from a biological perspective, they are typically queried independently for specific purposes, making it challenging to fully leverage the wealth of information they contain without effectively integrating them.

Knowledge graph (KG), a data structure depicting real-world entities and their relationships, has emerged as a favored method for integrating databases and deriving new insights from their connections, particularly in structuring biomedical knowledge for translational sciences (Nicholson and Greene, 2020; Fecho *et al*., 2022). KGs organize biological entities (e.g., genes, proteins, metabolic pathways, chemical compounds, and drugs) along with relationships between them into a coherent, interconnected network. By leveraging existing database knowledge integration, researchers can uncover hidden patterns and gain more insights into biology. For example, GenomicKB (Feng *et al*., 2023) unifies existing knowledge on the human genome, epigenome, transcriptome, and 4D nucleome within a large knowledge graph for exploring specific patterns between omics data. RTX-KG2 (Wood *et al*., 2022) combines data from 70 biomedical knowledge bases to create a KG on which powerful computational reasoning resource were developed for biomedicine (Ma *et al*., 2023). KG-COVID-19 (Reese *et al*., 2021) integrates COVID-19 data sources to expedite research in discovering new treatments for COVID-19. These success stories highlight the potential of employing knowledge graphs in the metagenomics field. However, there have been limited efforts directed towards such application in metagenomics. Currently, there is lack of a dedicated knowledge graph specifically designed to support effective metagenomic analysis. Joachimiak et al. (Joachimiak *et al*., 2021) built KG-Microbe using Natural Language Processing (NLP) techniques but did not include any biomedical information (e.g., diseases). Santangelo et al. (Santangelo *et al*., 2023) constructed MGMLink via gutMGene (Cheng *et al*., 2022) and PheKnowLator framework (Callahan *et al*., 2023) to evaluate mechanistic hypotheses about interactions between gut microbes and diseases. However, its application in metagenomics research is limited due to a reliance on a small number of gutMGene-described microbes and the lack of an efficient mapping method between new microbes and the knowledge graph. To date, there appears to be no existing metagenomics-specific, general purpose knowledge graph.

To resolve these limitations, we develop a new metagenomics-focused knowledge graph `MetagenomicKG`. It integrates the commonly-used taxonomic information (e.g., GTDB taxonomy and NCBI taxonomy) and other biomedical knowledge from several important resources, like KEGG, BV-BRC, and RTX-KG2, covering biological functions, systems, pathogenic microbes, phenotypic features, and diseases. `MetagenomicKG` is designed to enhance metagenomic data mining capabilities and explore the underlying relationships between microbes and diseases. To demonstrate its promising utility, we show a few use cases that might be useful in metagenomic research, which includes hypothesis generation and exploration for microbes and diseases, generation of sample-specific graph-based embeddings and pathogen predictions.

We envision the `MetagenomicKG` as a dynamic and versatile tool in the metagenomics field, offering a comprehensive and multi-modal reference framework tailored for metagenomic investigations. It is designed to support a wide spectrum of exploratory analyses and empower researchers to delve deeply into the complex interplay of microbial communities. Moreover, it acts as knowledge repository for machine learning endeavors, offering structured information that can drive the development of predictive models and innovative analytical tools. With the extensible nature of knowledge graphs, we anticipate the continual expansion of `MetagenomicKG` by integrating more established databases, thereby broadening its applicability to foster new discoveries and enhance our understanding of microbial life.

## 2 Graph Overview

Our `MetagenomicKG` integrates data from 7 sources: GTDB taxonomy (Parks *et al*., 2022), NCBI taxonomy (Schoch *et al*., 2020), KEGG (Kanehisa *et al*., 2017), RTX-KG2 (Wood *et al*., 2022), BV-BRC (Olson *et al*., 2023), MicroPhenoDB (Yao *et al*., 2020), and NCBI AMRFinderPlus Prediction (Feldgarden *et al*., 2021) (see Figure 1). `MetagenomicKG` is a directed multigraph with 1.25 million nodes and 56 million edges. The nodes are categorized into 14 distinct node types, which mostly align with the types of KEGG databases: Microbe, Phenotypic Feature, Disease, KEGG orthology (KO), Compound, Reaction, Drug, Glycan, Enzyme, Drug Group, Antimicrobial Resistance (AMR), Network, Pathway, Module. To better compare data and enable interoperability with other knowledge bases, we adhere to the semantic schemas of most existing biomedical knowledge graphs (Bizon *et al*., 2019; Reese *et al*., 2021; Wood *et al*., 2022; Morris *et al*., 2023) that map node types and edge types to the standard node category and predicates defined by the Biolink model (Unni *et al*., 2022). The Biolink model is a comprehensive, open-source framework that aims to standardize and facilitate the interoperability of diverse biological and translational science data across biomedical knowledge graphs. Table 1 shows all node types used in `MetagenomicKG` with their mapping to the Biolink entity category and their explanations. Figure 2 shows all Biolink edge predicates used in the graph and their distribution between different node types. Currently, the *biolink:genetically_associated_with, biolink:associated_with, biolink:subclass_of, biolink:superclass_of* are the top 4 predicates with the most edges in `MetagenomicKG`. These predicates mostly connect node types among Microbe, KEGG orthology (KO), Enzyme, Module, and Pathway.

**Table 1.**
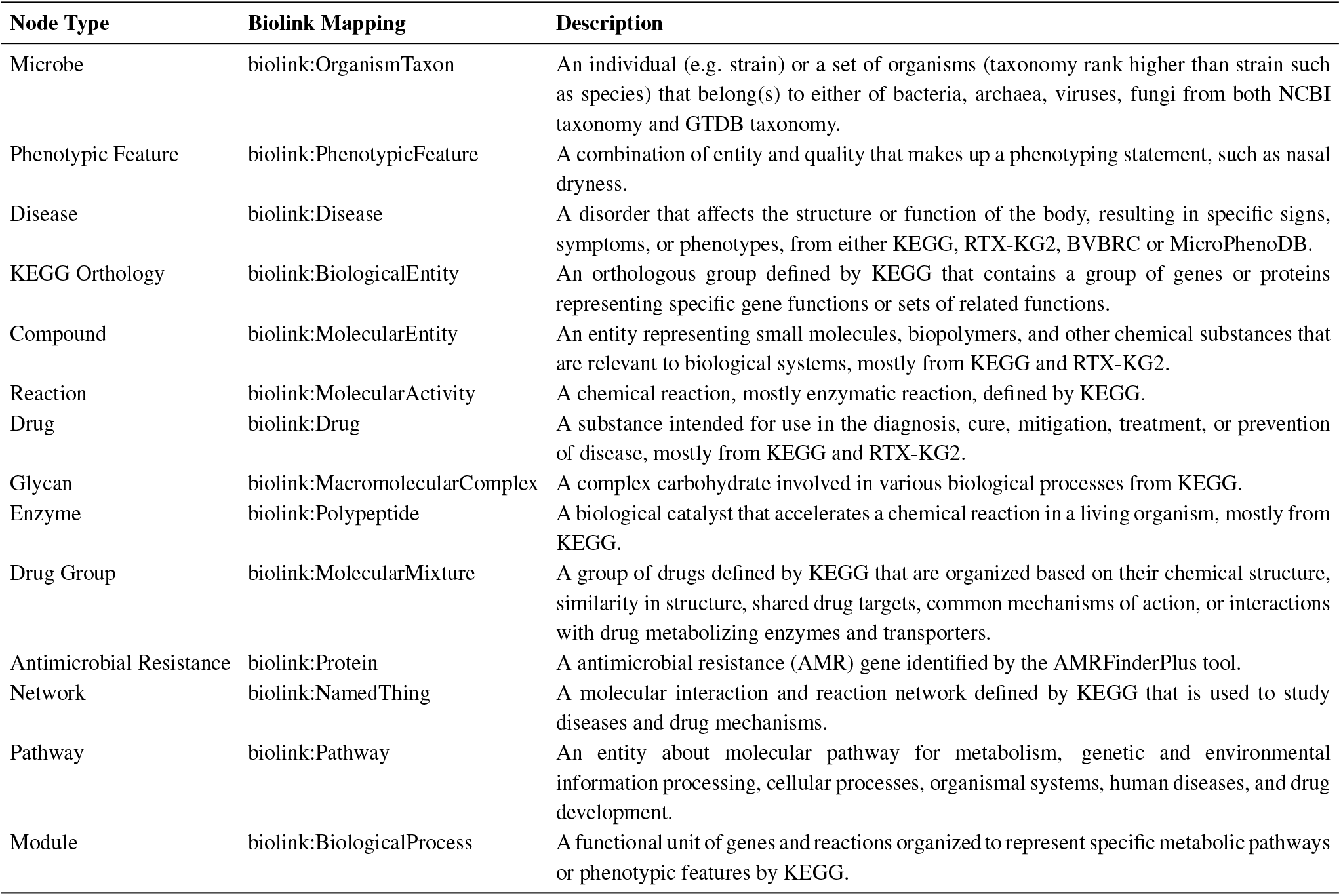
Description of node types used in MetagenomicKG.

**Fig. 1:**
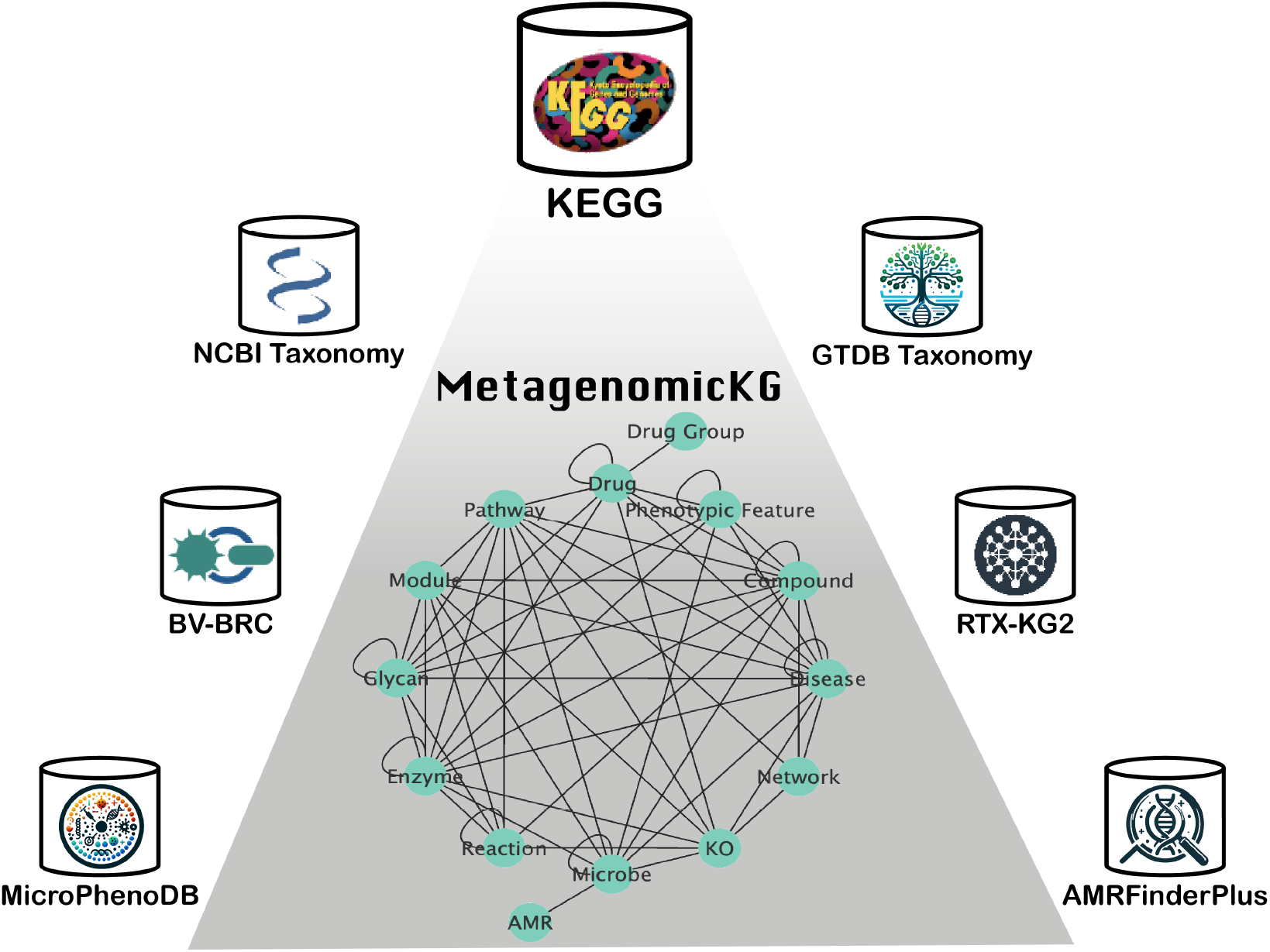
Overview of `MetagenomicKG`

**Fig. 2:**
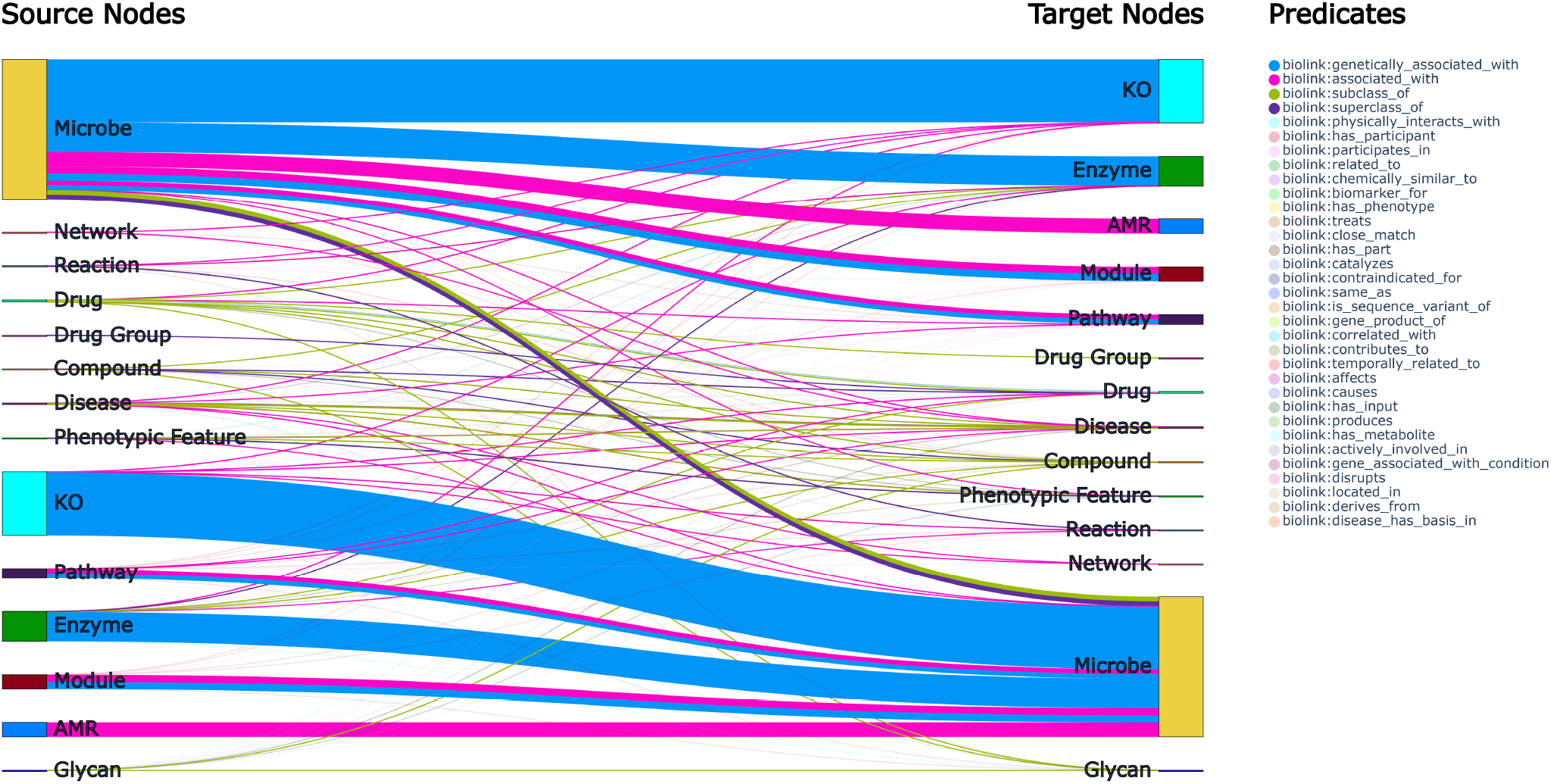
Sankey plot displaying the distribution of edge predicates between node types used in `MetagenomicKG`. The colored rectangles represent different node types while the colored flow lines correspond to different types of Biolink predicates used in the graph. The line thickness is proportional to the quantity of edge predicates between node types. To reduce graph complexity, we show the top 4 number of edge predicates with fully opaque, and the rest with 25% transparency.

### 2.1 Data Sources

Here we detail the seven data sources currently used in `MetagenomicKG`, which are categorized into four major categories. Each category is structured to be extensible, accommodating new data sources when available. This ensures that `MetagenomicKG` can adapt and grow, reflecting the continuous evolution of metagenomic data and research. Furthermore, we have enumerated several critical tools in establishing edges (connections) between biological entities within the graph, further enhancing the graph’s utility and interconnectivity.

#### 2.1.1 Taxonomy

*GTDB Taxonomy* – a standardized, genome-based taxonomy database for both bacteria and archaea built on a comprehensive phylogenetic approach. It incorporates large-scale, high-quality genomic data to establish a consistent and evolutionary framework for taxonomy classification, enabling a more accurate reflection of microbial diversity and evolutionary relationships. It has been widely adopted in the field of metagenomics (Nissen *et al*., 2021; Smith *et al*., 2022; Blanco-Míguez *et al*., 2023). We integrate its latest version (release214) into `MetagenomicKG`.

*NCBI Taxonomy* – the largest, comprehensive system for naming and classifying all sequenced organisms within the International Nucleotide Sequence Database Collaboration’s nucleotide and protein databases. It provides a standardized framework for the up-to-date classification and naming of viruses and fungi. We only utilize its viral and fungal taxonomy information (downloaded on 08/30/2023) in `MetagenomicKG`.

#### 2.1.2 Functional annotation

*KEGG* – an important database resource that stores comprehensive molecular and functional annotations based on genome sequences and other high-throughput data. It provides a wide range of databases, such as pathways, compounds, glycans, reactions, human diseases, drugs, drug group, functional orthologs, and their interconnections. The content of `MetagenomicKG` is mainly based on the data downloaded from its FTP server on 10/11/2023.

#### 2.1.3 Pathogen characterization

*BV-BRC* – an advanced research platform that emerged from the integration of the PATRIC (Gillespie *et al*., 2011), IRD (Zhang *et al*., 2017), and ViPR (Pickett *et al*., 2012) databases, providing abundant data for bacterial and viral pathogens. All bacterial and viral genomes along with their metadata were downloaded via the FTP server of BV-BRC on 06/07/2023. Only genomes with human as host and clearly associated diseases in the metadata are considered as pathogens and included into `MetagenomicKG`.

*MicroPhenoDB* – a database that maps the relationships between pathogenic microbes, their core genes, and human disease phenotypes. It was built through a manual review process and a calculation method that includes data from the Infectious Diseases Society of America (IDSA) guidelines and other manually curated data resources. Data from this database was downloaded on 05/30/2023.

#### 2.1.4 Existing Biomedical Knowledge Graphs

*RTX-KG2* – one of the largest open-source, Biolink model-based standardized Biomedical Knowledge Graphs that integrated extensive knowledge from 70 databases which include those curated by humans, derived through computational methods, or based on scholarly publications. The version 2.8.4 of RTX-KG2 that we used in `MetagenomicKG` contains around 6.8 million nodes and 45.36 million edges. Since this knowledge graph contains many node types that are not associated with metagenomics (e.g., “biolink:GeographicLocation” and “biolink:Device”), we only use the knowledge that is relevant for exploring relationships between microbes and diseases and hence extract from RTX-KG2 those nodes with type ‘biolink:Disease’ and ‘biolink:PhenotypicFeature’ and the edges adjacent to such nodes. We also exclude the RTX-KG2 edges that are only supported by the Semantic MEDLINE Database (SemMedDB) due to contradictory or erroneous relationships (Cong *et al*., 2018) caused by SemRep algorithm (Rindflesch and Fiszman, 2003).

#### 2.1.5 Bioinformatics tools

*GTDB-tk* – a taxonomic classification tool devised by the Genome Taxonomy Database (GTDB) team to provide standardized and objective taxonomic assignments to bacterial and archaeal genomes. It leverages a marker-gene-based reference tree, employs HMMER for gene alignment and pplacer for genomic placement within the reference tree. Subsequent classification is based on Relative Evolutionary Divergence (RED), Average Nucleotide

Identity (ANI), and the established phylogenetic positioning. We utilized GTDB-tk to harmonize genome fasta files from various databases, ensuring each new genome is accurately integrated into the appropriate node within our graph.

*KofamKOALA* – a tool designed by the Kyoto Encyclopedia of Genes and Genomes (KEGG) for the functional annotation of genome sequences. It employs a database of KEGG Orthology (KO) terms linked to specific metabolic pathways and functions, offering a precise understanding of protein roles within the cellular context. The tool uses HMMER-based searches to align input sequences against a set of curated reference profiles corresponding to the KO terms, ensuring the high-confidence prediction of protein functions. We applied KofamKOALA to annotate genomic sequences to build connections with the functional layer in our graph.

*NCBI AMRFinderPlus Prediction* – a tool developed by the National Center for Biotechnology Information (NCBI) to identify antimicrobial resistance (AMR) genes, resistance-associated point mutations, and other related genetic elements in bacterial genomes. It utilizes the BLASTX protein search against the NCBI Bacterial Antimicrobial Resistance Reference Gene Database for AMR protein identification. We applied this tool to the genome sequences of the databases mentioned above to predict their potential AMR genes.

### 2.2 Construction

The `MetagenomicKG` build process is automated via a parallel workflow framework, Snakemake. It uses a combination of Python and shell scripting. The information of all nodes and edges in the graph are stored in two separate files with a tabular format and imported into the Neo4j graph database for Cypher querying. Each node in the graph contains information including node id, node type, all possible names (“all_names” column), description, original data source (“knowledge_source” column), source links, synonyms, pathogen label (“is_pathogen” column). The identifier of each biological entity from the databases above is categorized into one of node types described in Table 1. It is assigned a new node id by combining the node type with an order number (e.g., 1, 2, …). The bacterial and archaeal genomes from the databases above were assigned to the closest GTDB identifier via GTDB-Tk (Chaumeil *et al*., 2020) if they lack RefSeq or GenBank assembly identifiers, with Average Nucleotide Identity (ANI) threshold *≥* 99.5%. Otherwise, other microbial genomes are merged based on their RefSeq and GenBank assembly identifiers. Different identifiers and names of the same biological entity are linked via the Unified Medical Language System (UMLS) search function^1^, Ontology mapping with Ontology Xref Service (OxO)^2^, as well as Node Synonymizer function provided by RTX-KG2 (Wood *et al*., 2022). Each edge contains the attributes of source node, target node, predicate description, and original data source (“knowledge source” column), where both source and target nodes use the new node id consistent with the one used in the node table. To increase the reliability of our nodes and edges, we can use the “knowledge source” and “links” attributes to trace back to their original data sources.

## 3 Applications of Metagenomics Knowledge Graph

In this section, we show that `MetagenomicKG` can have multiple applications for researchers in the field of metagenomics. We provide a few use cases to demonstrate how `MetagenomicKG` can be leveraged for metagenomics research.

### 3.1 Metagenomic Resource Integration for Hypothesis Generation and Exploration

The `MetagenomicKG` offers an integrated metagenomic resource with a reduction of redundancy. For example, more than 15,000 human pathogens have been recorded in the BV-BRC database but many of them are near-identical (sharing more than 99.5% ANI with other genomes). To better present the taxonomic relationships and reduce redundancy, we consolidated these genomes, as well as pathogens from the KEGG database, into 2,682 unique strain-level nodes that are attached to 405 species from GTDB. This aggregation facilitates a more efficient taxonomic representation and user-friendly exploration of underlying biological relationships between pathogenic genomes while preserving the scalability to include more pathogen annotations in the future.

To connect microbial genomes and diseases via the molecular functions and pathways, the `MetagenomicKG` connects genomes and their higher-rank taxonomic groups to KEGG functional layers, which can further link to other KEGG databases and thus unravel the functional mechanisms of pathogens and their roles in diseases. Although not all microbial genomes can be connected to KEGG databases, we can utilize shared proteins associated with Antimicrobial Resistance (AMR) genes and Virulence Factors (VF) to link genomes with unknown molecular mechanisms to those with known mechanisms. By linking `MetagenomicKG` with RTX-KG2, we further significantly broaden the scope from a focused microbiome perspective to encompass broader diseases and phenotypic features, thus enabling a wider range of practical applications.

With integration of the above resources, `MetagenomicKG` provides a more holistic approach to understanding and evaluating the relationships between pathogens, their functions, and potential drug targets. For example, Staphylococcus bacteria are usually harmless but cause serious infections that are hard to treat due to resistance to some antibiotics ^3^. We can illustrate this case in the `MetagenomicKG` in Figure 3. It shows a subgraph representing the association between AMR:streptomycin resistance protein (green node), Microbe:Staphylococcus aureus subsp. aureus ST398 (purple node), KO:K23587 (blue node), Pathway:map05150 (red node), Disease:Methicillin-resistant Staphylococcal aureus (MRSA) infection (brown node) and Drugs (orange node). This subgraph indicates that one of particular strain of Staphylococcus, known as Staphylococcus aureus subsp. aureus ST398 is capable of causing MRSA infection via the potential mechanism involving KO function unit of superantigen-like protein 5/11 and KO pathway of map05150. This pathogen has certain AMR protein CBA13544 and currently can be treated by several drugs. This illustration showcases the capability of the `MetagenomicKG` to facilitate the exploration of metagenomic data, which can be extended to tackle more complex analyses. For instance, another genome with similar characteristics, i.e. possesses crucial genes within the key pathway as well as AMR genes that counteract the drugs targeting this disease, is more likely to be identified as a pathogen.

**Fig. 3:**
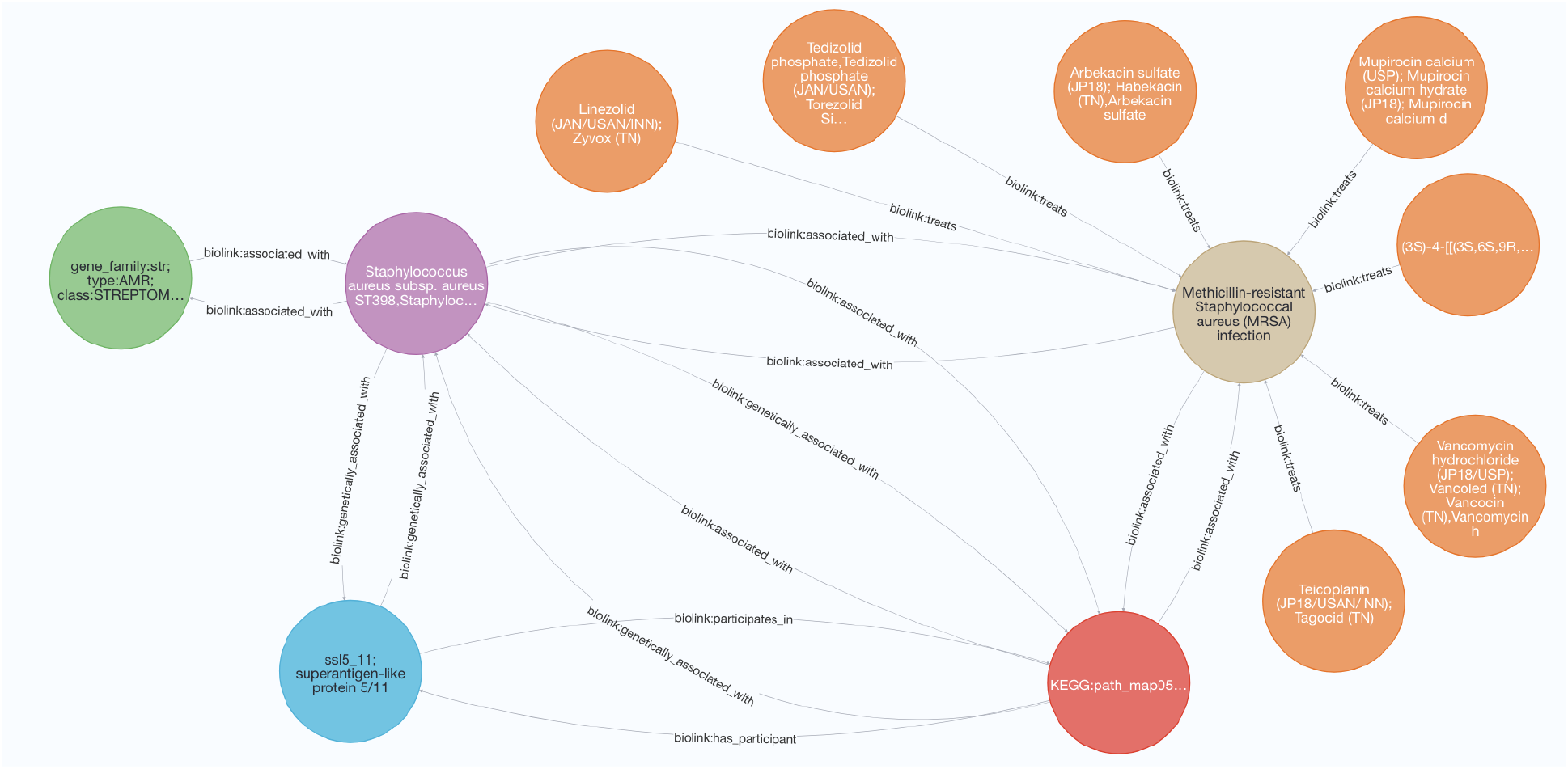
A subgraph representing the MRSA disease, its treatment, its causative pathogen, and relevant functional unit and pathway, as well as the known AMR protein.

### 3.2 Generation of Sample-specific Graph Embeddings

By utilizing graph embeddings to incorporate multiple metagenomic reference knowledge, `MetagenomicKG` with graph neural network techniques can present a novel approach for the exploratory data analysis of metagenomic samples. For example, while taxonomic and functional profiling are often used to characterize metagenomic communities, they don’t reveal the underlying biological relationships among microbes, metabolic pathways, and diseases. To better incorporate the existing biological knowledge into the representation of a metagenomic sample, we generate a sample-specific embedding (i.e. a numerical vector representation) using a modified version of the PageRank algorithm (Page *et al*., 1999), which has been demonstrated to efficiently integrate and embed knowledge from the knowledge graph and external data sources (Nelson *et al*., 2019). By utilizing this method, we can integrate taxonomic or functional profiling results with existing metagenomic knowledge from `MetagenomicKG` into sample-specific graph embeddings. The length of the embedding equals to the number of nodes in `MetagenomicKG`. Each value in the embedding corresponds to a node in the graph, with its value indicating the importance of that node for the given sample. These embeddings enable a range of downstream analyses, including differential analysis of important sample-related biological entities, classification analysis to predict disease status (Luo and Long, 2018; Huang *et al*., 2017), as well as network analysis to elucidate microbial community structures (Liu *et al*., 2021). They can facilitate the discovery of hidden patterns, connections, and insights that might be missed by conventional analysis techniques.

To illustrate this kind of approach, we use real data from the Human Microbiome Project (HMP) and BV-BRC databases. Taxonomic profiles, as well as sample-specific graph-based embeddings, were generated for 100 randomly selected samples from 5 body sites to perform clustering analysis. The results are visualized by Principal Coordinates Analysis (PCoA) in Figure 4 (See more method details in Section S2 in the supplementary material). The observation aligns with our expectation that diverse body sites have distinct microbial communities while closely related body sites (e.g., palatine tonsil, buccal mucosa, gingiva) may exhibit similar microbial compositions. Although both methods derive clear cluster boundary, but the graph-embedding-based approach on the right seems to capture more biological patterns, bringing samples from closely related body sites closer together. This may indicate, although not clearly demonstrated, the graph embeddings for metagenomic samples can help in prioritizing top features beyond taxonomic and functional entities for further analysis. This highlights the efficacy and promising utility of graph embeddings based on `MetagenomicKG` for metagenomic samples.

**Fig. 4:**
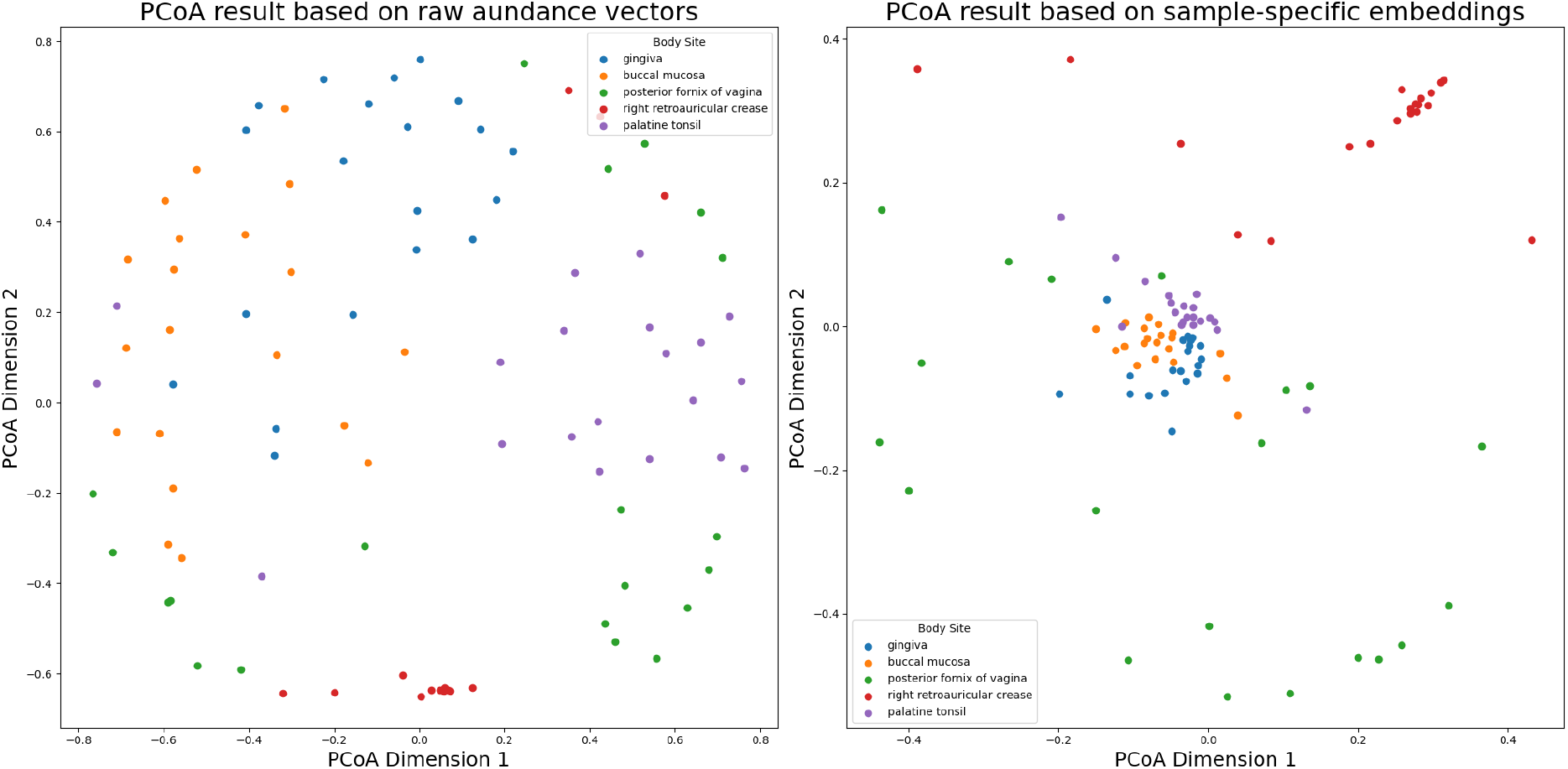
Comparison of PCoA visualizations for 100 random HMP samples between those using raw abundance generated by Sourmash and those using sample-specific embeddings generated by the personalized PageRank algorithm. The left figure is the PCoA result based on raw abundance while the right figure is the PCoA result based on the sample-specific embeddings. reliable human-related pathogens from KEGG and BV-BRC (see more details in section S3).

### 3.3 Pathogen Predictions

Detecting pathogenic organisms is crucial for clinical diagnostics, public health initiatives, and understanding environmental microbiomes (Ko *et al*., 2022; Wang *et al*., 2022; Yang *et al*., 2020). Traditional methods in metagenomics for pathogen detection rely on conserved gene families that are commonly shared among strains in the same species, such as 16S rRNA for bacteria and 18S rRNA for protozoa and fungi. These methods utilize deep amplicon sequencing (DAS) technique to amplify and identify taxonomic markers in pathogenic species (Miller *et al*., 2013). But these DAS-based methods require prior knowledge of potential pathogenic agents, causing the omission of pathogenic organisms with unknown markers. Unlike the DAS technique, shotgun sequencing can capture all nucleic acids present in a sample, enabling the detection of rare and unknown pathogens. Sequence-based models, such alignment-based methods ((Sf, 1990)) and k-mer-based methods ((Wood and Salzberg, 2014; Rosen *et al*., 2011)), have been popular in recent years. Machine learning methods have also evolved to predict pathogen genomes by learning sequence patterns or alignment status (Deneke *et al*., 2017; Bartoszewicz *et al*., 2020, 2021; Jiang *et al*., 2023). However, to our best knowledge, all existing machine learning approaches (e.g., PaBrBaG (Deneke *et al*., 2017), DeePac (Bartoszewicz *et al*., 2020)) rely on the sequence-based features from known pathogens for prediction. Such homology-based approaches can struggle to identify novel or distantly related pathogens. This leads to a major limitation for unknown pathogens when there are rare or no close reference genome exist in the training data. The `MetagenomicKG` can address this drawback by leveraging functional connections, taxonomic relationships, and additional biomedical connections to predict potential pathogenicity. Novel pathogens, which share limited sequence similarities with known reference database, can still be recognized through the graph structure which captures additional information about pathways, proteins, biological processes, etc. .

To demonstrate the advantage of `MetagenomicKG` in pathogen identification, we apply a simple graph neural network (GNN) to `MetagenomicKG` (see more details in section S3 in the supplementary material) and compare it with other state-of-the-art sequencing-based models in binary classification of pathogens. For ground-truth data, we collect a list of reliable human-related pathogens from KEGG and BV-BRC (see more details in section S3).

To show `MetagenomicKG`’s superiority in identifying pathogens lacking sequences similar to those in the training dataset, we strategically divided the ground-truth data into training, validation, and test sets based on clustering with average nucleotide identity (ANI) to avoid the training genomes similar to test genomes. We call this split method as “Missing Reference Split”. Specifically, we first utilize Sourmash (Pierce *et al*., 2019) to calculate the pairwise ANI scores of all ground-truth data. Then we cluster them according to the ANI scores. Figure 5 shows the hierarchical clustering of all ground-truth genomes in two dimensions based on 1 - ANI. We see that many positive/pathogenic ground-truth genomes are highly clustered together, indicating high sequence similarities. We intentionally pick a few clusters (e.g., cluster 15) with low similarity to the training set as the test set to simulate scenarios where we lack proper references for unknown pathogens. Table 2 shows the ground-truth data distribution after data split. As a comparison, we also train models using 10-fold Cross Validation random data split methods.

**Table 2.** Ground-truth data distribution for missing-reference data split.

| Data Type | Positive (pathogenic) | Negative (non-pathogenic) |
| --- | --- | --- |
| Training set | 2,413 | 2,616 |
| Validation set | 152 | 140 |
| Test set | 111 | 817 |

**Fig. 5:**
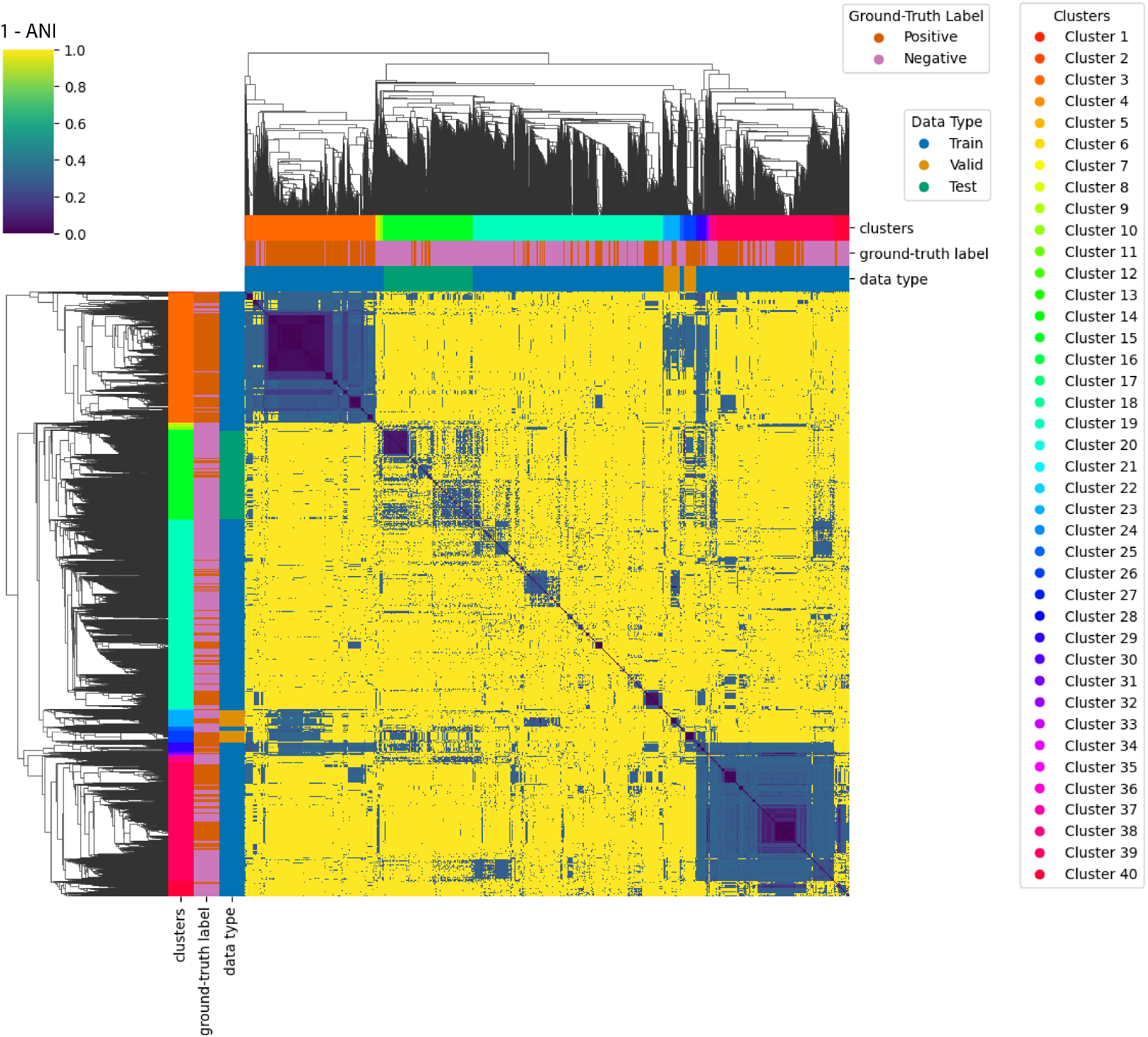
Heatmap plot with hierarchical clustering in two dimensions to show the sequence similarity of ground-truth genomes used in the “pathogen predictions” use case. The genomes are clustered into 40 groups based on 1 - ANI as distance. The three color bars correspond to ‘clusters’, ‘ground-truth label’, and ‘data type’ classifications, respectively. The ‘positive’ genomes are the ground-truth pathogenic microbes while the ‘negative’ genomes are the ground-truth non-pathogenic microbes.

We train the GNN model with `MetagenomicKG` and other sequence-based baseline models using the training data. Then, we evaluate their performance based on test data using 6 common metrics: Accuracy (ACC), True Positive Rate (TPR), True Negative Rate (TNR), Area Under the Receiver Operating Characteristic Curve (AUROC), Average Precision (AP), and F1 score (F1) (See more descriptions about these metrics in Section S3). Table 3 shows the performance comparison of GNN model with `MetagenomicKG` and all other baseline models in the task of pathogen predictions using two different data split methods. As shown in the table, when using the random split datasets, the baseline models can achieve good and comparable performance as the `MetagenomicKG`-based GNN model because of the reference genome homology to “unknown” pathogens (i.e. those in the test set). However, in the missing-reference data split, as expected, due to lack of homology to reference genomes, the sequence-based baseline models become significantly worse in identifying the “unseen” pathogens based on TPR, AP, and F1 metrics, while the `MetagenomicKG`-based GNN model still has robust performance. We highlight in particular around 20 percentage point improvement in TPR for the GNN with `MetagenomicKG` approach as compared to baseline models.

**Table 3.**
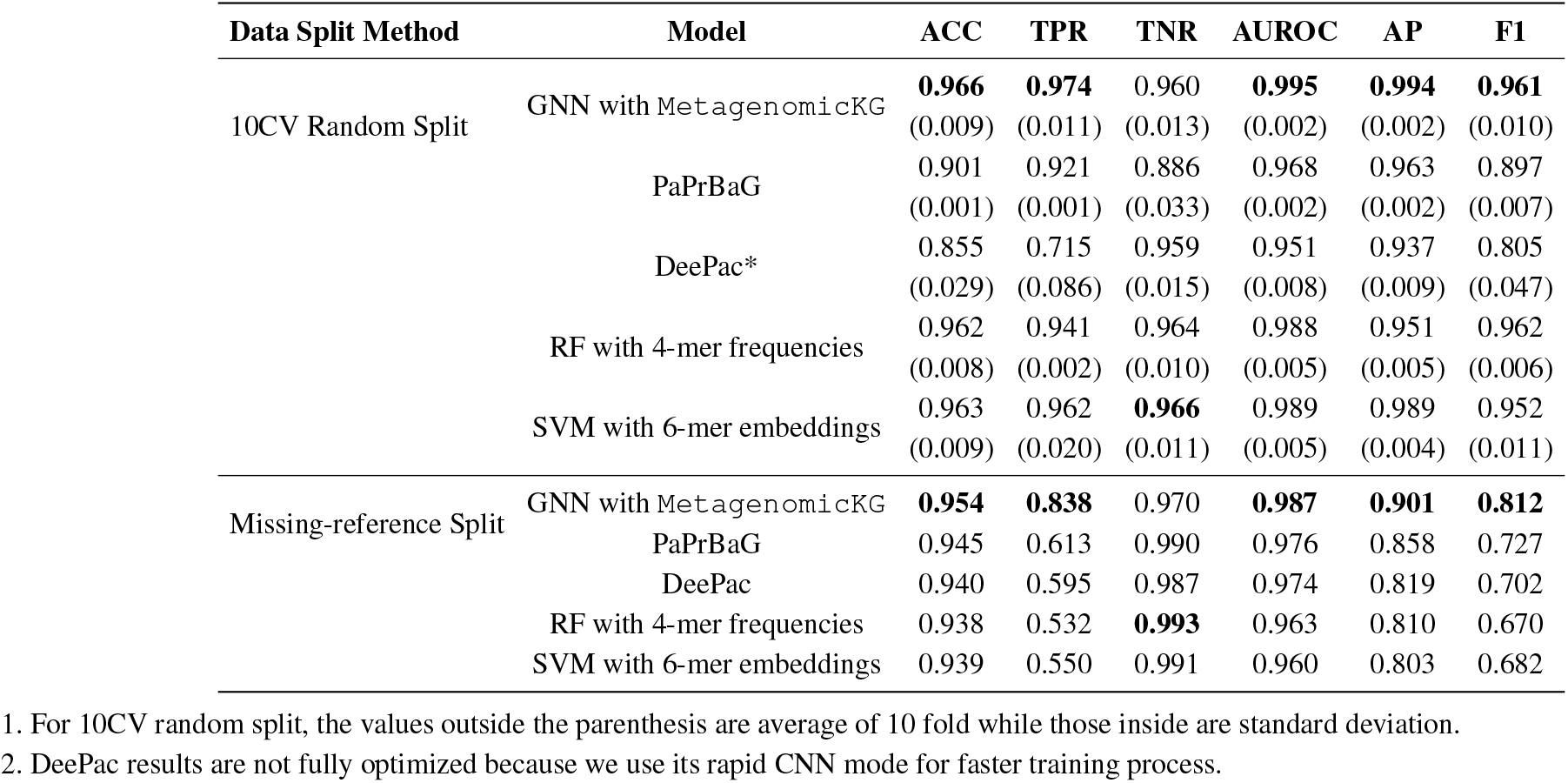
The performance comparison of pathogen prediction task between GNN with MetagenomicKG and different baseline models based on test dataset using random data split in 10 fold cross validation and missing-reference data split.

These results illustrate that the efficacy of sequence-based models depends on the similarity between captured features from training sequences and those of the “unseen” pathogens. However, there might be many pathogenic microbes in the “microbial dark matter” that are dissimilar to the known pathogens. In such cases, `MetagenomicKG` combined with GNN technique proves more effective than the sequence-based models for pathogen predictions.

## 4 Conclusion

This work introduce a novel metagenomics knowledge graph, `MetagenomicKG`, which integrates the commonly-used taxonomic taxonomy, functional annotations, pathogenic microbe information, and other biomedical knowledge from 7 relevant databases. This integration is aimed at enhancing our understanding of the relationships between microbes and diseases. To the best of our knowledge, this graph is the most comprehensive knowledge graph related to metagenomics to date. To demonstrate the application of `MetagenomicKG` in metagenomics research, we provide three different use cases. First, we show that using our `MetagenomicKG` can help understand the complex relationships among pathogens, AMR-associated proteins, KEGG-based function annotations, diseases, and relevant drugs. Researchers can utilize it to get new insights and evaluate their biological hypothesis. Second, we characterize metagenomic samples by embedding the taxonomic profiling with the topological structure of `MetagenomicKG`.

The embeddings enable better distinguishing the samples from different body sites when compared to taxonomic profiling results alone. Lastly, we find that combining graph neural network models with `MetagenomicKG` can improve the accuracy of pathogen identification compared with other sequence-based models, especially for those pathogenic microbes that have less sequence similarity with the known pathogens.

`MetagenomicKG` is mainly based on connections between a collection of biological databases in the Kyoto Encyclopedia of Genes and Genomes (KEGG), which contains comprehensive and high-quality functional and molecular information of biological systems. Using KEGG as basis, we further integrate further metagenomic knowledge (e.g., taxonomy, AMR, as well as relevant diseases and drugs). This can help researchers better understand the biological mechanisms underlying the impact of microbes on human health. Additionally, since we categorize all nodes and edges of `MetagenomicKG` using the standardized ontologies of Biolink Model, it facilitates easier extension and interoperability with other Biolink Model-based knowledge graphs, such as SPOKE (Morris *et al*., 2023), KG-COVID-19 (Reese *et al*., 2021), BioThings Explorer (Callaghan *et al*., 2023), mediKanren (ZHENG *et al*., 2020). We believe that the applications of `MetagenomicKG` in this work can serve as paradigms to indicate how knowledge graph can contribute to metagenomic research communities.

### Funding

Support for this work was provided by NCATS through the NIH award OT2TR003428. Any opinions expressed in this document do not necessarily reflect the views of NIH, NCATS, individual Translator team members or affiliated organizations and institutions.

## Footnotes

1 UMLS Search API: https://documentation.uts.nlm.nih.gov/rest/search/

2 OxO REST API: https://www.ebi.ac.uk/spot/oxo/docs/api

3 Centers for Disease Control and Prevention: Methicillin-resistant Staphylococcus aureus (MRSA)

