## Supplemental file for "MetagenomicKG: a knowledge graph for metagenomic applications"

#### S1 Resource integration

##### S1.1 Use GTDB-tk to assign taxonomic labels for BV-BRC and KEGG genomes

To unify taxonomic information for pathogen genomes from KEGG and BV-BRC, as well as to accommodate future resources, we employed GTDB-tk-2.3.2 [1] (bundled with GTDB version r214) for assigning taxonomic labels. Initially, we downloaded all genome FASTA files from the KEGG FTP server (which requires a license) and the BV-BRC FTP server. These FASTA files were then utilized as input for GTDB-tk. Subsequently, we obtained the default GTDB reference data by executing the bundled script "download-db.sh" within GTDB-tk, a process that may extend over several hours. Classification analysis was thereafter conducted using the command "gtdbtk classify_wf" with the default parameter and the "–skip_ani_screen" option.

Given that there are approximately 23,000 genomes to process, we leveraged GNU parallel to expedite this procedure. For detailed execution scripts and further instructions, please refer to our GitHub repository.

##### S1.2 Identify AMR and VF genes by AMRFinderPlus

We employed AMRFinderPlus [2] to detect antimicrobial resistance (AMR) genes and virulence factors (VF) within bacterial genomes. The National Center for Biotechnology Information (NCBI) maintains a meticulously curated reference database for AMR and VF, which we incorporated as gene nodes within our knowledge graph. The identification of these genes is represented as edges in the graph.

AMRFinderPlus can analyze both pure DNA sequences and DNA sequences accompanied by protein sequences (faa files) and protein annotation files (gff files), with a preference for the latter approach. We initially utilized the "NCBI datasets" tool to download all relevant genome files, when available. Subsequently, AMRFinderPlus was run with default parameters based on these input files. For detailed execution scripts and further instructions, please refer to our GitHub repository.

#### S2 Usecase 2: Generation of Sample-specific Graph Embeddings

For reproducible analysis, we've included the raw metadata and bash scripts in our GitHub repository.

##### S2.1 Data Selection

All human metagenomic samples analyzed in this study were obtained from the Human Microbiome Project (HMP) data portal [3], utilizing the search parameters "fastq" for format and "WGS raw seq set" for type. We further refined our selection to include only samples from healthy individuals, narrowing the total to 1,864 samples. To ensure sufficient depth and coverage, samples with a file size of less than 200 MB were excluded, resulting in a final dataset of 1,674 samples. Although we possessed the complete data matrix, we randomly selected a subsample to succinctly showcase our results.

We first identified 7 largest groups in the dataset: gingiva, dorsum of tongue, buccal mucosa, external naris, posterior fornix of vagina, right retroauricular crease, and palatine tonsil. The "dorsum of tongue" group was excluded as we already have selected samples from the mouth region, which are similar; the "external naris" group was also omitted due to its proximity to, yet slight differentiation from, the "right retroauricular crease" group. For each remaining 5 groups, 20 samples were randomly selected using the command "shuf –random-source= < (yes 33)" on the metadata of all files.

Taxonomic profiling was then conducted using Sourmash gather [1] with a $k$-mer size of 31, based on the GTDB r214 reference. The relative abundance data from each sample were merged for both Principal Coordinates Analysis (PCoA) and the generation of graph embeddings.

##### S2.2 Embedding Generation and Clustering Analysis

To generate graph-based embeddings for 100 randomly selected metagenomic samples, we first utilized Sourmash gather function to obtain taxonomic abundance of potential microbes, also known as taxonomic profiling, for each one. Then, these results were transformed into an abundance table, with each row representing microbial genomes and each column representing a sample. The table values indicate the relative abundance of specific microbes in each sample (if the sample lacks of a specific microbes, its abundance is set to 0). Then, we normalized the sample-specific relative abundance in each column and used each normalized column vector as a personalized vector for the PageRank algorithm for the corresponding sample. The personalized PageRank algorithm was implemented by the NetworkX python package [2]. The converged output vector is the sample-specific graph-based embeddings.

To perform clustering analysis using raw relative abundance vectors and the sample-specific graph-based embeddings, we utilized Principal Coordinates Analysis (PCoA) to do dimension reduction and visualized them in a 2D figure. Instead of using the entire sample-specific graph embeddings, for each of 140 samples, we extracted the top 1,000 significantly relevant nodes based on the embedding values. Then, we merged these top 1,000 nodes from all samples and used their corresponding embedding values as a subset of sample-specific embeddings for t-SNE visualization.

#### S3 Usecase 3: Pathogen Predictions

For reproducibility analysis, we've included the analytical scripts in our GitHub repository.

##### S3.1 Ground-truth Data Preparation for Pathogen Predictions

To prepare ground-truth data for training models in the use case of pathogen predictions described in the main text, we used the KEGG microbial organisms labeled as "Human pathogen" (e.g. 'T00329') and the BV-BRC microbial genomes whose host is human and is associated with a concrete disease. After filtering out duplicate genomes, we finally obtained 2,676 known human pathogens as positive ground-truth data. We obtained non-pathogenic microbial genomes by profiling healthy samples, in which 15 different body sites are covered, from the Human Microbiome Project (HMP) [4]. We used YACHT [5], a state-of-the-art metagenomic detection tool, to identify potential organisms in these samples. The parameters for kmer size were set at 31 and minimum coverage at 0.01. We used the latest version (rs214) representative genomes of GTDB as the reference database. After aggregating all detected microbial organisms and filtering out duplicates, we obtained a total of 3,573 non-pathogenic organisms as negative ground-truth data.

##### S3.2 GNN model

In this experiment, We used a GNN model structure that consists of 1 linear transformation layer, followed by 3 GraphSAGE layers [6] with mean aggregator, and 1 binary transformation layer. This model structure is implemented using Deep Graph Library (DGL) [7]. For parameter settings, we set the embedding size of all linear layers and GraphSAGE layers to 512, the number of neighbor nodes sampled in each GraphSAGE layer to 100, the learning rate to 0.001. We used a maximum of 500 epochs to train the GNN model with early stopping and a batch size of 256. The model was trained on a NVIDIA A100 80GB. To incorporate other information from `MetagenomicKG`, we concatenated the name and node type of each node to generate embeddings using a biomedical language model called PubMedBERT [8], which was reduced to 200 dimensions using Principal Component Analysis (PCA) and then served as input for training the proposed GNN model.

##### S3.3 Baseline models for comparison

In our study, we evaluated models based on 2 categories: contig/read classifiers such as PaPrBaG [9] and DeePac [10] and whole genome classifier such as BacPaCS [11] and DCiPatho [12]. PaPrBaG learns from thousands of sequence features, including motifs, codon usage, k-mers, and spaced k-mers, from input contig sequences. It aggregates its

[1] https://sourmash.readthedocs.io/en/latest/classifying-signatures.html#projecting-abundances-in-sourmash-gather
[2] NetworkX: https://networkx.org/

predictions at the genome level through a majority vote strategy. Similarly, DeePac, a read-level classification tool that utilizes DNA one-hot embeddings from fragmented genomes as input for a reverse complement neural network, makes fragment-level predictions and then use a majority vote. We followed their documents with the standard data process for data preparation, model training and prediction. It should be noted that DeePaC becomes extremely computationally intensive when training on thousands of genomes. To accommodate this, we opted for its rapid CNN mode to expedite data processing, rather than the more sensitive LSTM mode, which may lead to a relatively lower performance.

Regarding the genome-level prediction, BacPaCS, which focuses on protein sequences, may not be a good fit here. In order to do comprehensive comparison, we also utilized two additional common classifiers: a k-mer-frequency-based random forest classifier and a k-mer-embedding-based SVM classifier, both leveraging 4-to-6 k-mer frequencies. Therefore, DCiPatho, another tool that utilizes frequency features of 3-to-7 k-mers for whole-genome prediction was not included in comparison. For the random forest classifier, we created a k-mer frequency table for each genome as input for training. In contrast, for the SVM classifier, we transformed k-mer profiles of each genome into embeddings using the word2vec package, and then averaging these embeddings as the basis for SVM classifier training.

##### S3.4 Description of Evaluation Metrics used in Pathogen Predictions

We used the following 6 metrics to evaluate the performance of the GNN model and baseline models used in the application pathogen predictions:

Accuracy (ACC) is the proportion of correct predictions in the model classification, calculated as:

$$\text{ACC} = \frac{\text{Number of correct classifications}}{\text{Total number of ground-truth labels}} \tag{1}$$

True Positive Rate (TPR), also known as sensitivity, is the proportion of accurate model predictions for genomes that have positive ground-truth labels, calculated as:

$$\text{TPR} = \frac{\text{Number of correct positive classifications}}{\text{Total number of positive ground-truth labels}} \tag{2}$$

True Negative Rate (TNR), also known as specificity, is the proportion of accurate model predictions for genomes that have negative ground-truth labels, calculated as:

$$\text{TNR} = \frac{\text{Number of correct negative classifications}}{\text{Total number of negative ground-truth labels}} \tag{3}$$

Area Under the Receiver Operating Characteristic Curve (AUROC) is the area under the curve plotted with the true positive rate (TPR) against the false positive rate (FPR) across various thresholds. The equations of TPR and FPR are respectively defined as:

$$\text{TPR} = \frac{\text{True Positive}}{\text{True Positive} + \text{False Negative}} \qquad \text{FPR} = \frac{\text{False Positive}}{\text{False Positive} + \text{True Negative}} \tag{4}$$

where $TruePositives(TP)$ is the number of positive ground-truth genomes that are predicted to be pathogenic by the model. $FalsePositive(FP)$ is the number of negative ground-truth genomes that are incorrectly predicted to be pathogenic by the model. $FalseNegative(FN)$ is the number of positive ground-truth genomes that are incorrectly predicted to be non-pathogenic by the model. $TrueNegative(TN)$ is the number of negative ground-truth genomes that are correctly predicted to be non-pathogenic by the model.

Average Precision (AP) is defined as the average of the precision values at different recall levels, measuring the area under the precision-recall curve, calculated as:

$$\text{AP} = \sum_n (R_n - R_{n-1}) P_n \tag{5}$$

where $P_n$ and $R_n$ are the precision and recall at the nth threshold.

F1 score (F1) is defined as a harmonic mean of the precision and recall, calculated as:

$$\text{F1} = \frac{2 * TP}{2 * TP + FP + FN} \tag{6}$$

### References


[1] Pierre-Alain Chaumeil, Aaron J Mussig, Philip Hugenholtz, and Donovan H Parks. Gtdb-tk: a toolkit to classify genomes with the genome taxonomy database, 2020.

[2] M Feldgarden, V Brover, N Gonzalez-Escalona, JG Frye, J Haendiges, DH Haft, M Hoffmann, JB Pettengill, AB Prasad, GE Tillman, et al. Amrfinderplus and the reference gene catalog facilitate examination of the genomic links among antimicrobial resistance, stress response, and virulence. sci rep 11: 12728, 2021.

[3] Peter J Turnbaugh, Ruth E Ley, Micah Hamady, Claire M Fraser-Liggett, Rob Knight, and Jeffrey I Gordon. The human microbiome project. *Nature*, 449(7164):804–810, 2007.

[4] HMP Integrative. The integrative human microbiome project: dynamic analysis of microbiome-host omics profiles during periods of human health and disease. *Cell host & microbe*, 16(3):276–289, 2014.

[5] David Koslicki, Stephen White, Chunyu Ma, and Alexei Novikov. YACHT: an ANI-based statistical test to detect microbial presence/absence in a metagenomic sample. *Bioinformatics*, 40(2):btae047, 01 2024.

[6] Will Hamilton, Zhitao Ying, and Jure Leskovec. Inductive representation learning on large graphs. *Advances in neural information processing systems*, 30, 2017.

[7] Minjie Yu Wang. Deep graph library: Towards efficient and scalable deep learning on graphs. In *ICLR workshop on representation learning on graphs and manifolds*, 2019.
